# A bibliometric analysis of Soil remediation Based on Massive research literature data During 1988-2018

**DOI:** 10.1101/689018

**Authors:** Ya Hu, Jichang Han, Zenghui Sun, Huanyuan Wang, Xiang Liu, Hui Kong

**Affiliations:** Shaanxi Provincial Land Engineering Construction Group, Xi’an 710075, China, Xi’an 710075, China; Key Laboratory of Degraded and Unused Land Consolidation Engineering, the Ministry of Land and Resources, Xi’an 710075, China; Institute of Land Engineering and Technology, Shaanxi Provincial Land Engineering Construction Group., Xi’an 710075, China; Shaanxi Provincial Land Consolidation Engineering Technology Research Center, Xi’an 710075, China; Xi’an Jiaotong University,710048, China

## Abstract

Soil is an important part of the ecosystem with significant roles that help human population sustain. Research on prevention and remediation of soil pollution has been carried out when 1985. This study analyzed the 1988–2018 soil remediation dataset in the Web of Science database by bibliometric methods to illustrate the current research trends and hot topics of quantitative analysis and soil remediation in the world. To further identify the major soil contamination topics, we employed social network analysis. The results indicate that the field of soil remediation has entered a stage of rapid progress. The United States has a strong overall strength with the largest number of published articles and larger impact. China ranks second. We identified Journal of hazardous materials as the most influential journal and Chinese academy of sciences as the most influential institution. Academic cooperation showed an increasing trend at the author, institutional, and national levels with an average level of cooperation of 3.57, 1.66, and 1.16, respectively. However, the growth rate of cooperation at the national level is relatively low. In addition, the frequency and co-word analyses of keywords revealed the important research topics. “heavy metals”, “PAH”, “bioremediation”, “Phytoremediation” and “Electrokinetic remediation” were identified as the hot topics. The findings of this study will help researchers understand the status of soil remediation as well as provide guidance for future research.

## Introduction

Soil is an important dependence of human survival. contaminated soil which polluted by heavy metals, agricultural inputs and solid waste, deteriorates the environment and restricts human development[1-3]. Soil contamination and remediation are global problems that have attracted the attention of governments and researchers[4]. In order to protect soil and prevent further deterioration, various studies have been conducted on remediation of contaminated soil. Many soil remediation technologies have been developed during the past few decades on different aspects such as chemistry, biology, agroecology, and electrodynamics[5,6]. During this period, new research ideas, methods, and means were introduced, and the remediation technology system was improved. At the same time, the intersection of discipline such as soil, engineering, chemistry and new materials promoted the rapid progress in soil remediation research[7]. However, the future of soil remediation technologies is uncertain and the need for multidisciplinary research is high. To gain research progress in soil remediation, we should focus on the key processes in soil remediation and break through the bottlenecks. We should explore new remediation technologies and perform a quantitative analysis of the relevant information in the field.

Bibliometrics can explores structures, characteristics, and laws of science and technology [8]. We used bibliometric method to analyze current research and the development trends in the field of soil remediation including total number of articles, countries’ performances, productive journals, performances of authors and institutios, citation, and extent of academic collaboration. This work will fill the gap in the field of soil remediation. Using frequency analysis and co-occurrence analysis of high-frequency keywords will help other researchers grasp the essence of advanced topics in this field. Based on the analysis, potential limitations and directions were derived to provide guidance to plan and implement future research.

## Materials and methods

### Data source

Data used in this study were taken from the Web of Science (WOS) core collection including Science Citation Index Expanded, Social Sciences Citation Index, Conference Proceedings Citation Index-Science, Conference Proceedings Citation Index-Social Science & Humanities, and Emerging Sources Citation Index. We searched the title, abstract, and keywords of 13891 published articles using 1988–2018 as the time phase, “soil remediation” as the keyword, and “subject” as the field.

The search date is January 18, 2019. The WOS derived document records included titles, authors, abstracts, and keywords. These indicators were analyzed using BibExcel, Ucinet and VOSviewer.

A general statistical analysis was performed on national distribution, journals, topics, authors, institutions, and citations. In addition, impact factor, academic cooperation, and national comprehensive strength were used to reflect the current academic impact of a country and of an author. Research and development in the field of soil remediation was analyzed, which will help researchers and policy makers attain an overall understanding of the subject.

### Impact factor

As the most commonly used assessment tool in bibliometrics, impact factor helps assess the merit of journals, authors, institutions, and countries [9].We collected the impact factors from the ISI Journal Citation Reports to evaluate the quality of the journals.

### Academic cooperation

Cooperation in scientific research is improving at all levels and in all areas, and this is a common indicator to measure closeness of collaboration in scientific research [10]. The indicators at all levels (author, institution, and nation) were used to calculate the degree of academic cooperation in the field of soil remediation. Equations used for calculation are as follows:

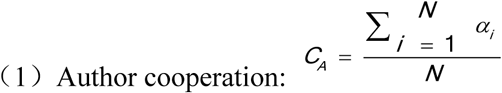

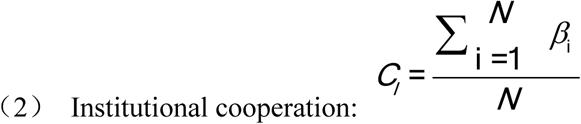

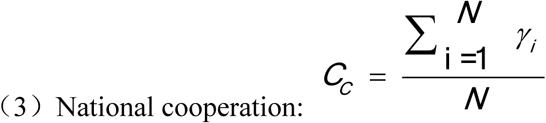

Where, G_A_, G_I_, and G_C_ represent the degree of cooperation by author, institution, and country, respectively; α_i_, β_i_, and γ_i_ represent the number of authors, institutions, and countries contributing to each paper, respectively; N represents the total number of articles in the field.

### Academic scale

Academic influence and academic competitiveness reflect a country’s comprehensive research strength. Four indicators were selected to assess national comprehensive research strength: (1) total number of articles (2) total citations (3) number of authors, and (4) number of research institutions. By calculating the standard scores of these four indicators, the combined score of each country was obtained, and the formulae used for calculation are as follows:

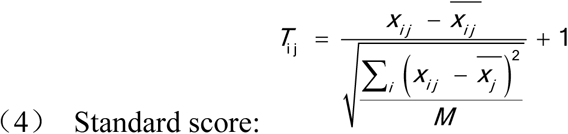

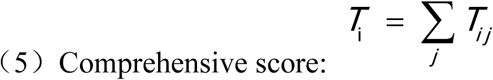

Where, T_ij_ represents the standard score of indicator j in country i; x_ij_ represents the original score of indicator j in country i; 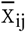 represents its average score; T_i_ represents the sum of the standard scores in country i; and M represents the number of countries.

## Results

### Contribution of country

The number of articles in a specific area is an important indicator to assess development trend. Analysis revealed that a total of 13,891 journal articles were retrieved from 148 countries and regions including England, Scotland, Wales, and Northern Ireland; China included only mainland China, and Hong Kong, Macao, and Taiwan were analyzed as separate regions. Different colors represent the number of articles in different geographical regions(**Fig 1**), the darker the color, the more the number of articles. The articles on soil remediation were mainly from the United States, China, Spain, and Canada. Research in this field was also prominent in Korea, Italy, and Australia.

**Figure 1.**
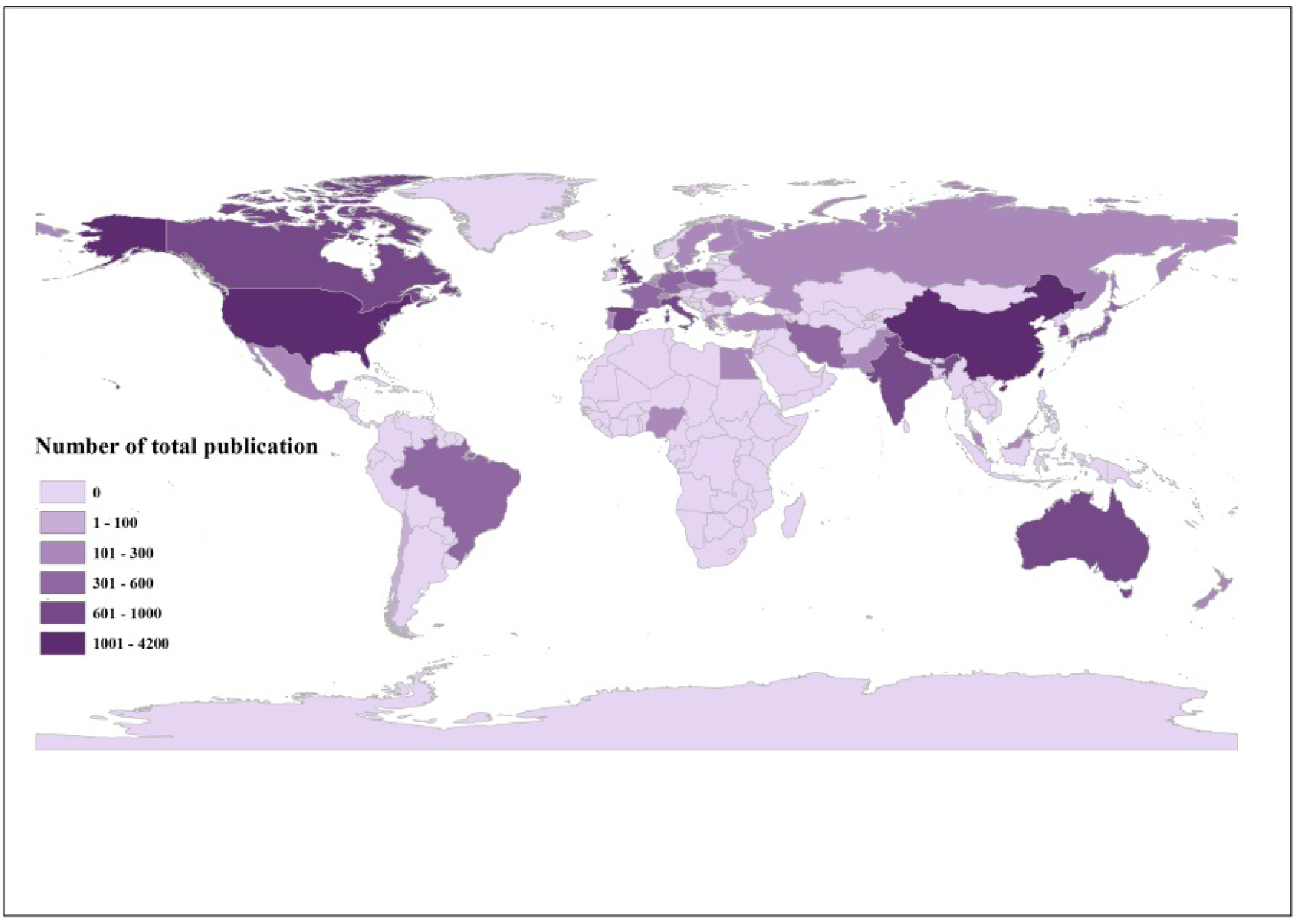
World map showing the distribution of research articles

The number of articles in the soil remediation field has grown rapidly over time (**Fig 2**), and the growth happened in three phases. Only a few developed countries such as the United States and Canada published few articles in the early beginning phase (1988 to 1998). In order to improve the quality of cultivated soil, healthy human living environment, many countries began to pay attention and study soil issues in the stable development phase (1998 to 2008). China’s soil remediation research is developing rapidly in the rapid growth phase (2008 to 2018).

**Figure 2.**
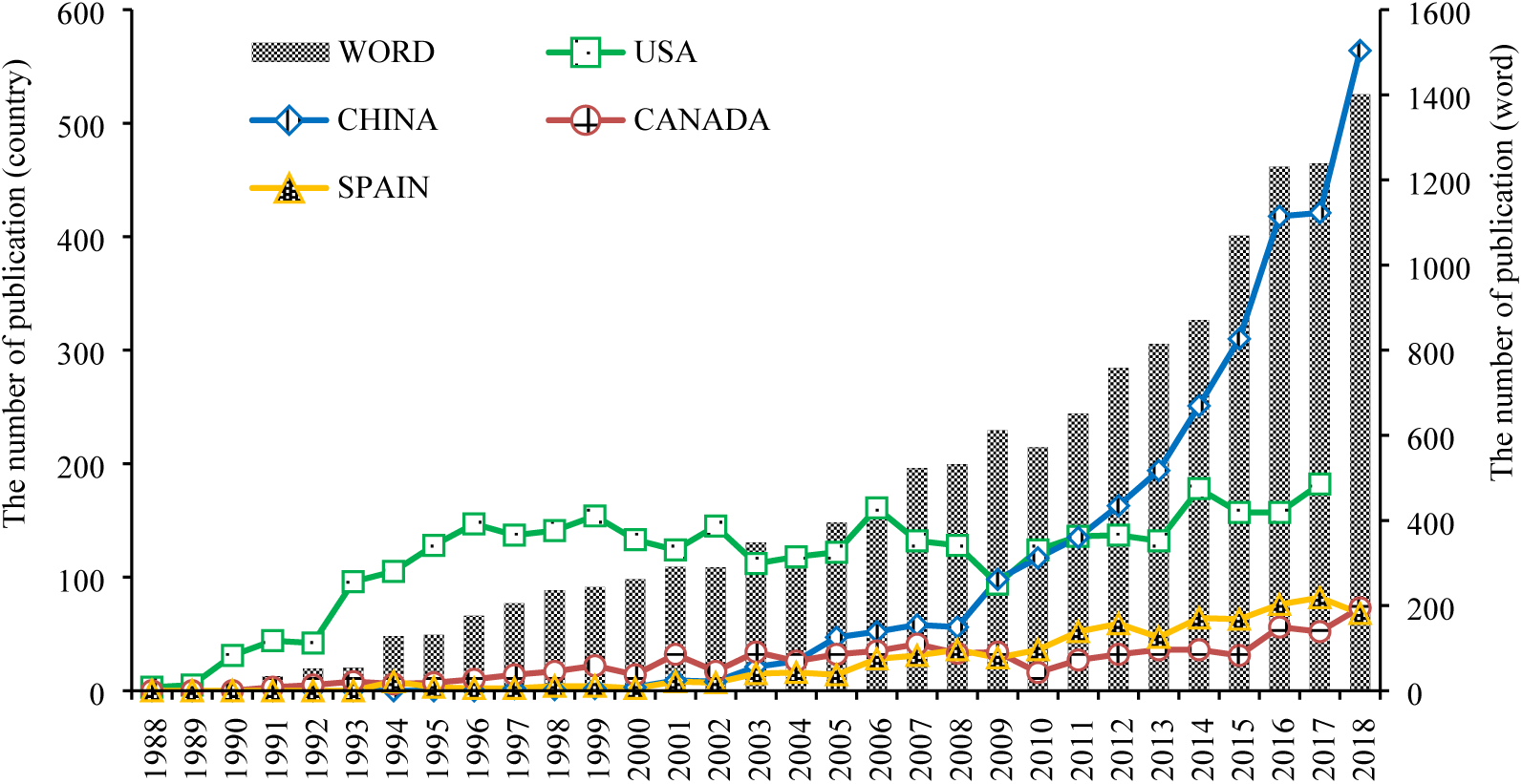
Distribution of major publishing countries

### Productive institutions

6637 institutions contributed to the soil remediation field. Among developing countries, four institutions among the 10 most published research institutions were in China (**Table 1**), the Chinese Academy of Sciences and the University of Chinese Academy of Sciences ranked first and third. Other research institutions were from developed countries, four from the United States and two from France. The Chinese Academy of Sciences contributed 706 articles (56.54% of the total number of Chinese articles). This indicates that Chinese Academy of Sciences is in a leading position in the field of soil remediation.

**Table 1.** Productive institutions during 1988-2018.

| RANK | Institution | Country | TP | TPRW(%) |
| --- | --- | --- | --- | --- |
| 1 | Chinese Academy of Sciences | China | 706 | 5.1 |
| 2 | United states department of energy doe | USA | 369 | 2.7 |
| 3 | University of chinese academy of sciences | China | 234 | 1.7 |
| 4 | Centre national de la recherche scientifique | France | 230 | 1.7 |
| 5 | Consejo Superior de Investigaciones Cientificas | France | 227 | 1.6 |
| 6 | university of california system | USA | 206 | 1.5 |
| 7 | Institute of soil science | China | 204 | 1.5 |
| 8 | Zhejiang University | China | 179 | 1.3 |
| 9 | United states department of agriculture | USA | 177 | 1.3 |
| 10 | State University System of Florida | USA | 175 | 1.3 |

Note: TP is the number of total articles; TPRW(%) is the ratio of the number of journal's publications in which institution to the total number of articles.

### Productive authors

32534 authors contributed to the soil remediation field. The authors with the most recent articles were from Denmark, and the productive authors were from other developed countries such as South Korea, United States, Australia, and Spain among the top 10 authors in the field of soil remediation(**Table 2**).

**Table 2.** Productive authors during 1988-2018.

| Rank | Authors | Country | TP | TC | CPP |
| --- | --- | --- | --- | --- | --- |
| 1 | Ottosen LM | Denmark | 82 | 1574 | 19.2 |
| 2 | Beak K | Korea | 70 | 1091 | 15.59 |
| 3 | Reedy KR | USA | 70 | 2347 | 33.53 |
| 4 | Naidu R | Australia | 66 | 1032 | 15.64 |
| 5 | Canizares P | Spain | 59 | 766 | 13.15 |
| 6 | Rodrigo MA | Spain | 59 | 736 | 12.47 |
| 7 | Tsang DCW | Hong Kong | 57 | 981 | 11.21 |
| 8 | Luo YM | China | 56 | 1173 | 20.95 |
| 9 | Lestan D | Slovenia | 52 | 1485 | 28.56 |
| 10 | OK YS | Korea | 52 | 1732 | 33.31 |

Note: TP is the number of total articles; TC is the number of total citations; CPP is citations per publication.

The most productive author was Ottosen LM (Denmark) has contributed to 82 articles. He mainly studied the use of electrodialysis technology and the use of electricity to deal with copper, lead, zinc, and chromium in industrial and mining fields [11,12]. Reddy K R (United States) was the highest cited author, whose articles has been cited 2347 times and mainly about electrodynamic remediation of heavy metals in soil [13]. Baek K (South Korea)was the most productive author in Asia with 70 articles. He analyzed the effect of electrolyte regulation of acidic and alkaline solutions on electroremediation of contaminated soil [14].

### Journals performance

1423 academic journals retrieved in the soil remediation field. These articles related to environmental science, soil contamination and botany. Journal of hazardous materials had 822 articles (5.9%)was the most published journal (**Table 3**).Chemosphere had 758 articles (5.5%) was the second most published journal. Environmental science and technology ranked fourth among all publications in all journals, however it has the largest impact factor (6.653) and had the most citations (27199).

**Table 3.** Top fifteen productive journals during 1998-2018.

| Rank | Journal | TP | TPR(%) | IF | TC |
| --- | --- | --- | --- | --- | --- |
| 1 | Journal of hazardous materials | 822 | 5.92 | 6.434 | 24491 |
| 2 | Chemosphere | 758 | 5.46 | 4.427 | 21607 |
| 3 | Environmental science and pollution research | 529 | 3.81 | 2.8 | 3985 |
| 4 | Environmental science technology | 482 | 3.47 | 6.653 | 27199 |
| 5 | Science of the total environment | 400 | 2.88 | 4.61 | 9032 |
| 6 | Water air and soil pollution | 369 | 2.66 | 1.769 | 5574 |
| 7 | Environmental pollution | 303 | 2.18 | 4.358 | 12079 |
| 8 | International journal of phytoremediation | 239 | 1.70 | 4.005 | 3371 |
| 9 | Journal of environmental management | 223 | 1.61 | 1.886 | 2592 |
| 10 | Journal of contaminant hydrology | 206 | 1.48 | 2.405 | 7230 |

Note: TP is the number of total articles; TPR(%) is the ratio of the number of journal's publications
to the total number of articles; IF is impact factor in 2018; TC is the number of total citations.

Among the top five journals, Journal of hazardous materials maintained a leading position in the number of articles from 2008 to 2013 (**Fig 3**). However, the number of articles in Environmental science and pollution research showed a rapid growth with an average growth rate of 30% per year after 2013. Science of the total environment showed short-term fluctuations but maintained high growth rate after 2013. Meanwhile, Journals related to this field from other journals showed relatively low growth rates.

**Figure 3.**
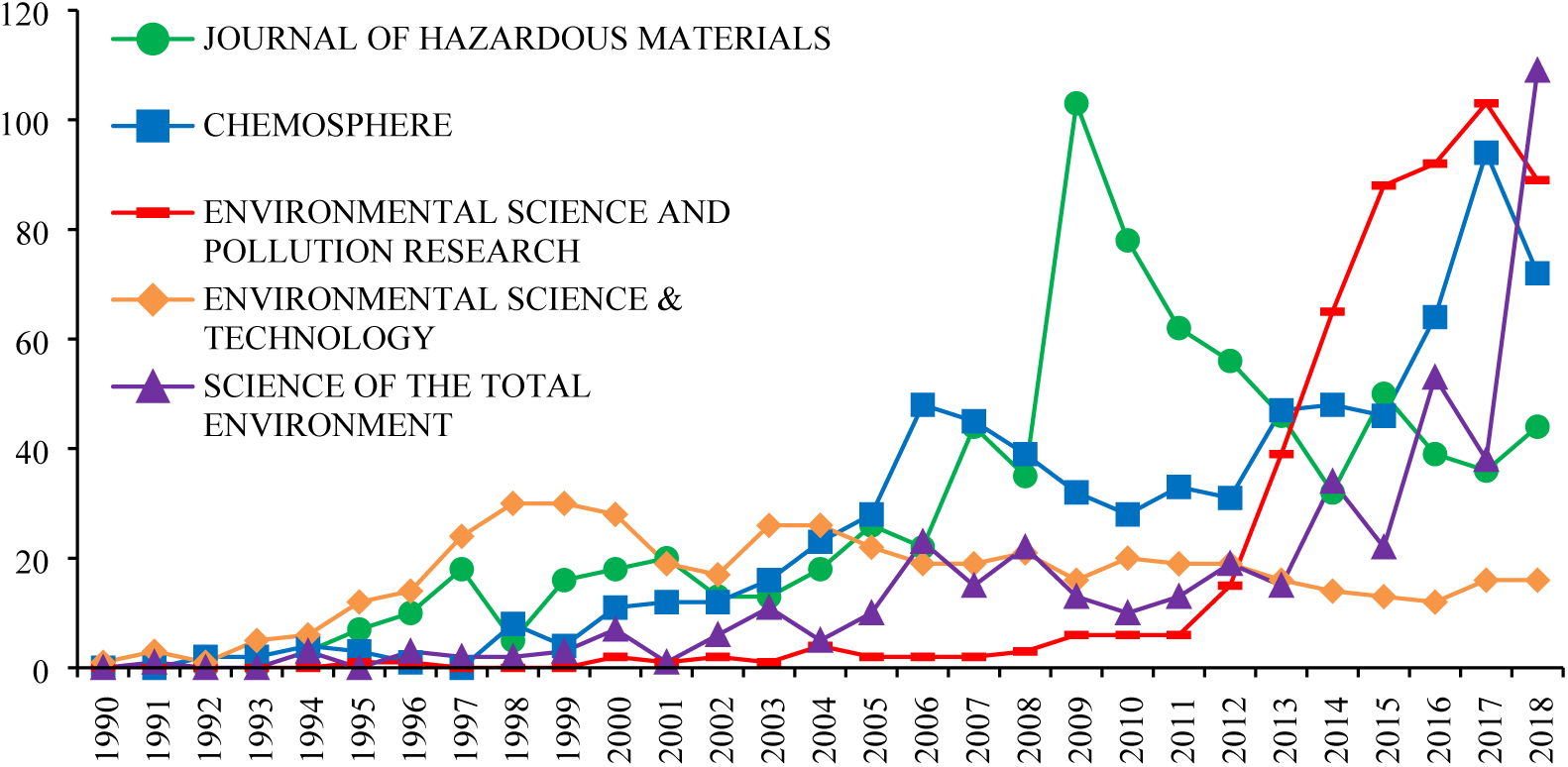
Trend of top five journals

### Subjects performance

Lists Environmental sciences ecology was the most popular subject with 8550 articles (61.5%) followed by Engineering and Water resources (29.1% and 11.6%) among the top 10 subjects closely related to the field of soil remediation(**Table 4**). Articles in this field focused on natural science subjects especially environmental science, ecology, geology, and meteorology and few social science subjects such as business and economics. Some domain-specified subjects including chemistry, agriculture, plants, and toxicology also published numerous articles because of their sensitivity to soil remediation.

**Table 4.** Distribution of subjects during 1988-2018.

| Rank | Subject | TP | TPR(%) |
| --- | --- | --- | --- |
| 1 | Environmental sciences ecology | 8550 | 61.5% |
| 2 | Engineering | 1043 | 29.1% |
| 3 | Water resources | 1612 | 11.6% |
| 4 | Agriculture | 1348 | 9.7% |
| 5 | Chemistry | 1176 | 8.4% |
| 6 | Geology | 863 | 6.2% |
| 7 | Biotechnology applied microbiology | 781 | 5.6% |
| 8 | Meteorology atmospheric sciences | 434 | 3.1% |
| 9 | Science technology other topics | 433 | 3.1% |
| 10 | Toxicology | 404 | 2.9% |

Note: TP is the number of total articles; TPR(%) is the ratio of the number of journal's publications to the total number of articles.

### Academic collaboration

The degree of academic cooperation reflects the degree of academic research in scientific research in this field. The degree of cooperation between authors, institutions, and countries was calculated using formulae (1), (2), and (3). Authors, institutions, and countries had cooperation levels of 3.57, 1.66, and 1.16, respectively, which indicate that 3.57 authors, 1.66 institutions, and 1.16 countries contributed to each article (**Fig 4**).The level of cooperation constantly improved, and the authors’ cooperation was Significant (3.57). Growth in national cooperation was slow due to the soil contamination problems are more concentrated in individual countries.

**Figure 4.**
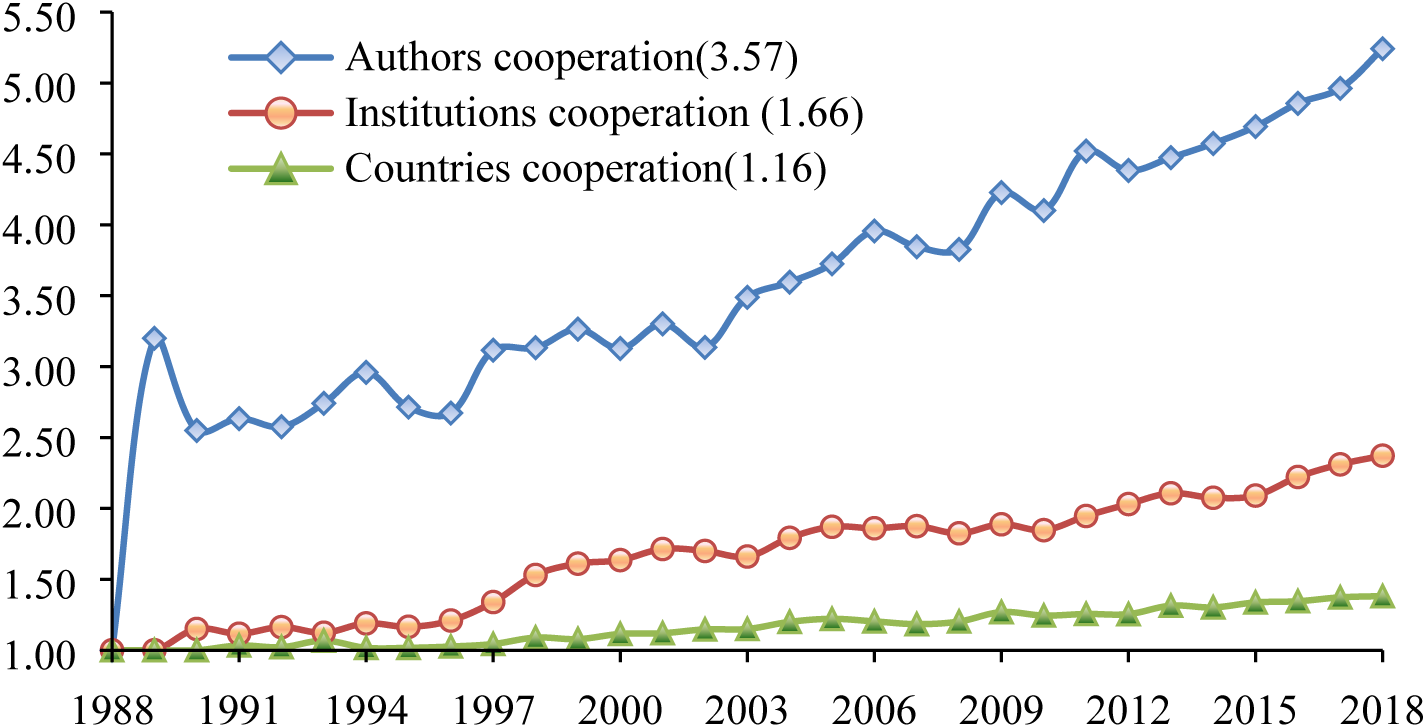
Academic cooperation during 1988-2018

## Research hot points

### Keyword clustering and frequency analysis

Keywords reflect the aim of research and summarize the key contents of the paper. We analyzed 1039 keywords used in 13891 articles through BibExcel. The keyword “remediation”demonstrated the highest frequency of occurrence (4372 times). According to the formula 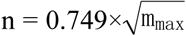 where m_max_=4372. In this study, n=50 implies that the keywords which are cited more than 50 times sre the core of the soil remediation field. We further classified 63 core keywords into 5 categories and labeled the number of occurrence for each keyword (**Table 5**).

**Table 5.**
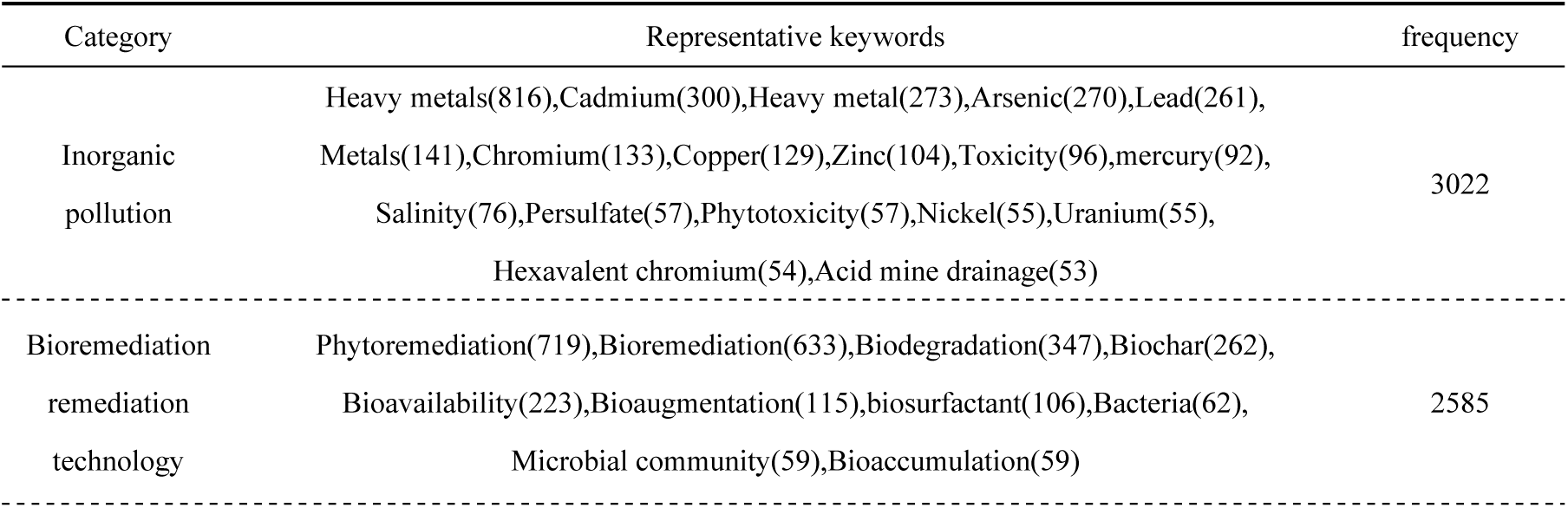
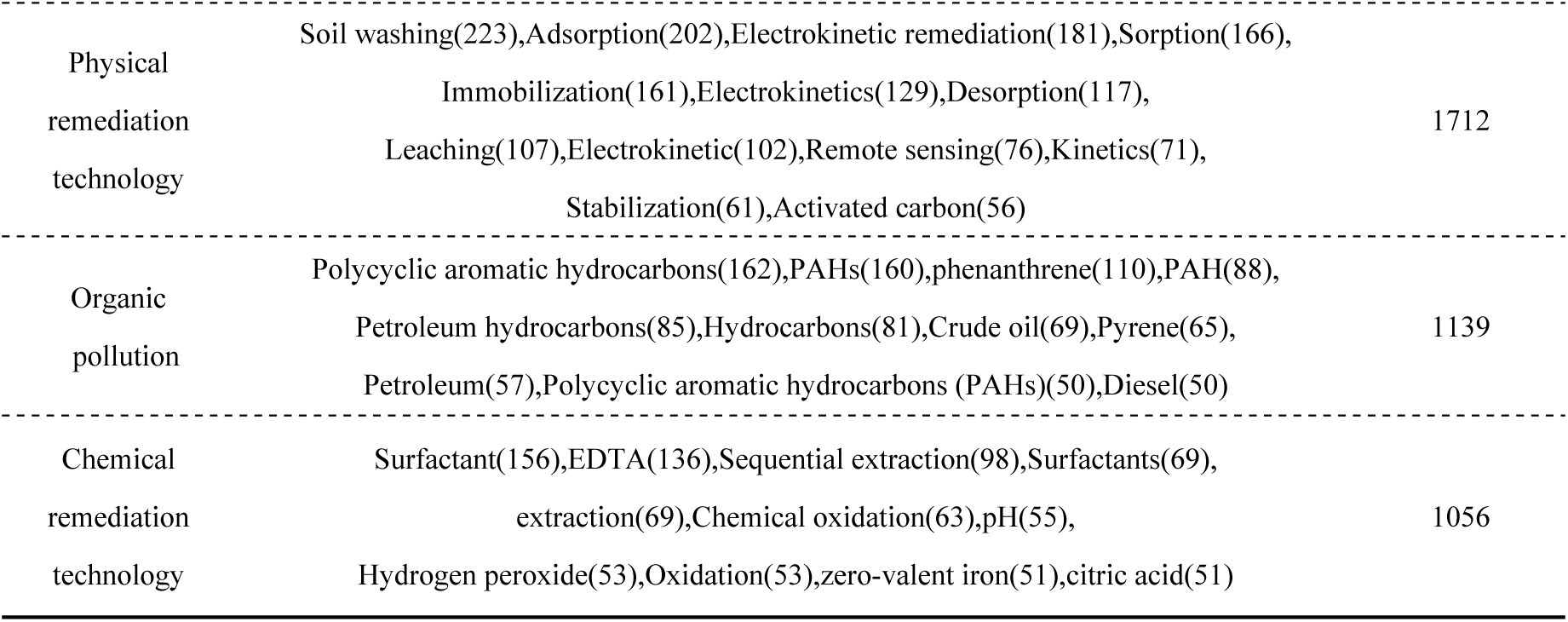
Frequency of keywords in soil remediation during 1988-2018.

#### Inorganic pollution

Heavy metals entered the soil with the rapid development of the global economy. Anthropogenic activities such as mining, industrial production, agriculture, and transportation are the major sources of heavy metals in soil. They cannot be completely removed from the soil by degradation and has caused soil contamination problems in many countries [15]. So “heavy metal” was a prominent keyword in soil contamination. Cadmium, arsenic, lead, chromium, and zinc have been the focus of remediation research followed by toxicity and salinization [16].

#### Bioremediation remediation technology

Bioremediation uses plants, animals, and microorganisms in the soil to absorb, degrade, and transform the soil Contaminants. “Phytoremediation” and “bioremediation” have been the focus of scholars after 2010 [17]. Bioremediation degrades Contaminants in situ at a low cost of remediation and with no secondary pollution. Due to the limitations of single remediation techniques, joint remediation techniques such as co-bioremediation remediation, physical-biological remediation, and chemical-biological remediation have been considered by some scholars.

#### Physical remediation technology

In situ soil washing, adsorption, immobilization, and other electric methods are the most studied physical methods [18]. Soil washing remediation mainly improves extraction efficiency by finding new eluents. Electrokinetic remediation was the most concerned chemical remediation technologys. In addition to new electrode technology which was represented by electrolyte optimization and approaching anode, Combined technology was represented by electric-permeable wall began to appeared and developed rapidly.

#### Organic pollution

Currently, more attention is paid polycyclic aromatic hydrocarbons, tocrude oils and petroleum hydrocarbons in organic pollution [19]. Pollution form Atrazine, chlorpyrifos and dichlorodiphenyltrichloroethane were maily in China, and explosive chemicals were mainly in the USA. With the increasing consumption of US military explosives, greatly studied researched on the soil organic pollution of military bases. China, India, Spain, and Canada fourced on the topic of petroleum, crude oil, and Petroleum hydrocarbons.

#### Chemical remediation technology

The main research in the field of chemical remediation technology was based on the chemical properties of pollutants or contaminated media. This method changed the chemical properties by the application of various chemical reagents, and separates the pollutants. Surfactants solubilize and elute soil contaminants, and EDTA complexes with the salts of heavy metals and increase the transport rate of heavy metals in soil. Thus, surfactants and EDTA were the most concerned chemical remediation technologys [4].

### Co-occurrence and network analysis of keywords

Co-occurrence analysis of high frequency keywords was performed using VOSviewer. The common keyword “remediation” was deleted in case it affected the display of other keywords.

Each keyword is represented by a circle in the visualization result of keyword average time distribution(**Fig 5**). The diameter of the circle and the size of the label indicate the appearance of keywords. The bigger the circle, the more the number of occurrences of the keyword [20].The distance between the circles indicates the proximity of the two keywords. The color of the circle represents the average publication year of the keywords. Lines represent co-occurrence links between two keywords. The thicker the line between two keywords, the more frequently they appear together.

**Figure 5.**
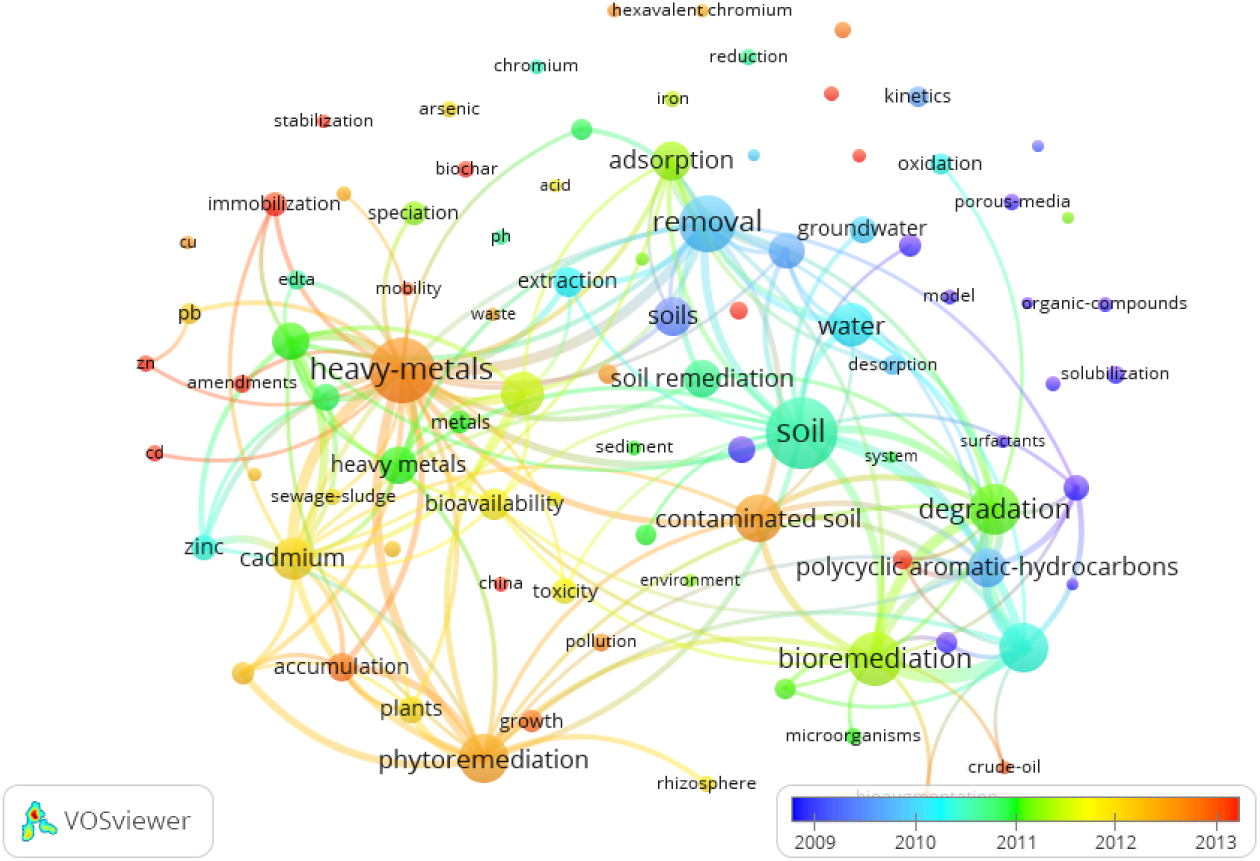
Co-occurrence map of keywords in the field of soil remediation

Heavy metals have more links to other keywords. This means that heavy metals have the maximum connection and reflects its central position in research. Large-scale keywords such as “bioremediation”, “phytoremediation”, and “removal” play an important role in the research network. Among all these keywords, the number of connections between “heavy metals”, “cadmium”, and “removal” was the most. This indicates their relevance and these three keywords or two of them usually appear together in the same research and most scholars focused on these issues. Another keyword group including “phytoremediation”, “bioremediation”, and “contaminated soil” also showed strong correlation, which indicates their relevance in soil remediation research.

Change in color indicates the trend in hot topics in this field. Blue represents the keywords that were released before 2009, such as “removal”, “grandwater”, “polycyclic aromatic-hydrocarbons” and “phenanthrene”. These words focused on PAH pollution and the migration mechanism of contaminants in soil. Green represents the keywords around 2011. Increase in soil contamination threatens the living environment and food security. Therefore, scholars have been paying more attention to soil contamination and remediation. The research techniques used were mainly bioremediation technologies including phytoremediation. As an important source of soil contamination, heavy metals continue to receive widespread attention. Studies have focused on “biochar”, “China”, “sewage”, and other specific topics and regions since 2013, rather than abstract and macro themes. These trends indicate that major research in the field of soil remediation is shifting from a contamination mechanism to technology application.

## Conclusions

Based on the WOS core database, the overall research development in the field of soil remediation from 1988 to 2018 was analyzed using bibliometric methods.

Soil remediation field has developed rapidly since 2008. The continuous increase in number of articles indicates that soil remediation is receiving increasing attention. At the national level, the United States had high overall strength with the largest number of articles and greater academic influence. As a representative of developing countries, China’s institutions and authors performed well by contributing more number of articles. The top five most published journals contributed 21.5% of all articles in the field in which Journal of hazardous materials was the most published journal. In addition, soil remediation included multidisciplinary fields, and environmental science ecology, engineering, and water resources were the top three subjects that published the most articles. Ottosen LM (Denmark), Reddy KR (the United States) and Baek K (Korea) were the authors with the more number published articles in this field. Academic cooperation showed an increasing trend at the author, institutional, and national levels with an average level of cooperation of 3.57, 1.66, and 1.16, respectively.

Cluster analysis and frequency analysis of the keywords indicate that the hot topics in this field were heavy metals, phytoremediation, bioremediation, electrodynamics, cadmium, leaching, solidification, and polycyclic aromatic hydrocarbons. According to the co-word analysis, “heavy metals” keyword had the maximum connection among all keywords and often appeared simultaneously with other keywords reflecting heavy metals as the core issue in the field. Trends in the hot topics in this field were discussed through the analysis of the keywords in published works. We found that research focus is shifting from the mechanism of pollutant transport in contaminated soils to the application of comprehensive repair technologies such as bioremediation technology and electric remediation. Meanwhile, research focused on specific micro-level issues and quantitative analysis in areas and regions. This study will help researchers understand the development trends in soil remediation research and provide guidance for future research.

However, this study only explores the overall theme changes of soil remediation research in the past 30 years from the time dimension. In order to further analyze the mature and declining process of prominent keywords, it is necessary to further classify and analyze the keywords on a case-by-case basis and summarize the development and decline process of keywords.

## Acknowledgement

This work was Supported by the Scientific Research Item of Shaanxi Provincial Land Engineering Construction Group (DJNY-2019-26),and the Fund Project of Shaanxi Key Laboratory of Land Consolidation (2018-TD02).

## References

1. Chen, H., Teng, Y., Lu, S., Wang, Y., Wang, J. (2015). Contamination features and health risk of soil heavy metals in China. Science of the Total Environment, 512-513,143–153.

2. Ni, Z., Wang, S. (2015). Economic development influences on sediment-bound nitrogen and phosphorus accumulation of lakes in China. Environmental Science and Pollution Research, 22, 18561–18573.

3. Khatri, N., Tyagi, S. (2015). Influences of natural and anthropogenic factors on surface and groundwater quality in rural and urban areas. Frontiers in Life Science, 8, 23–39.

4. Pietro, P., Falciglia, D. M., Vagliasindi, F. G.A. (2016). Removal of mercury from marine sediments by the combined application of a biodegradable non-ionic surfactant and complexing agent in enhanced-electrokinetic treatment. Electrochimica Acta, 222, 1569–1577.

5. Guarino, C., Sciarrillo, R. (2017). Effectiveness of in situ application of an Integrated Phytoremediation System (IPS) by adding a selected blend of rhizosphere microbes to heavily multi-contaminated soils. Ecological Engineering, 99, 70–82.

6. Ait Ahmed, O., Derriche, Z., Kameche, M., Bahmani, A., Souli, H., Dubujet, P., Fleureau, J.M. (2015). Electro-Remediation of Lead Contaminated Kaolinite: An Electro-Kinetic Treatment. Chemical Engineering and Processing: Process Intensification, S0255270115301537.

7. Mao, G., Shi, T., Zhang, S., Crittenden, J., Guo, S., Du, H. (2018). Bibliometric analysis of insights into soil remediation. Journal of Soils and Sediments, 18, 2520–2534.

8. Garousi, V., Mantyla, M. V. (2016). Citations, research topics and active countries in software engineering: A bibliometrics study. Computer Science Review, 19, 56–77.

9. Paulus, F. M., Rademacher, L., Schäfer, T., Müller-Pinzler, L., Krach, S. (2015). Journal Impact Factor Shapes Scientists’ Reward Signal in the Prospect of Publication. Plos One, 10, 0142537.

10. Wei, Y. M., Yuan, X. C., Wu, G., Yang, L. X. (2014) Climate Change Risk Assessment:A bibliometric Analysis Based on Web of Science. Bulletin of National Natural Science Foundation of China, 5, 347–356.

11. Jakobsen, M. R., Fritt-Rasmussen, J., Nielsen, S., Ottosen, L. M. (2004). Electrodialytic removal of cadmium from wastewater sludge. Journal of Hazardous Materials, 106, 127–132.

12. Jensen, P. E., Ottosen, L. M., Pedersen, A. J., Speciation Of Pb In Industrially Polluted Soils. Water, Air, and Soil Pollution, 107, 359–382.

13. Reddy, K. R., Parupudi, U. S., (1997). Devulapalli S N, et al. Effects of soil composition on the removal of chromium by electrokinetics. Journal of Hazardous Materials, 55, 135–158.

14. Baek, K., Kim, D. H., Park, S. W., Ryu, B. G., Bajargal, T., Yang, J. (2009). Electrolyte conditioning-enhanced electrokinetic remediation of arsenic-contaminated mine tailing. Journal of Hazardous Materials, 161, 457–462.

15. Chen, M., Xu, P., Zeng, G., Yang, C., Huang, D., Zhang, J. (2015). Bioremediation of soils contaminated with polycyclic aromatic hydrocarbons, petroleum, pesticides, chlorophenols and heavy metals by composting: Applications, microbes and future research needs. Biotechnology Advances, 33, 745–755.

16. Toth, G., Hermann, T., Da Silva, M. R, Montanarella, L. (2016). Heavy metals in agricultural soils of the European Union with implications for food safety. Environment International, 88, 299–309.

17. Hussain, I.,Aleti, G., Naidu, R., Puschenreiter, M. (2018). Microbe and plant assisted-remediation of organic xenobiotics and its enhancement by genetically modified organisms and recombinant technology: A review. Science of The Total Environment, 628, 1582–1599.

18. Derakhshan, N. Z, Jung, M. C., Kim, K. H. (2017). Remediation of soils contaminated with heavy metals with an emphasis on immobilization technology. Environmental Geochemistry and Health, 40, 927–953

19. Ivshina, I., Kostina, L., Krivoruchko, A. Kuyukina, M., Peshkurc, T., Anderson, P. et al. (2016). Removal of polycyclic aromatic hydrocarbons in soil spiked with model mixtures of petroleum hydrocarbons and heterocycles using biosurfactants from Rhodococcusruber IEGM Journal of Hazardous Materials, S0304389416302229.

20. Wang, Z., Zhao, Y., Wang, B. (2018). A bibliometric analysis of climate change adaptation based on massive research literature data. Journal of Cleaner Production, 199, 1072–1082.

